# Patient derived models of bladder cancer amplify tumor specific gene expression compared to surgical specimen while maintaining gene expression of molecular subtype and epithelial mesenchymal transition markers

**DOI:** 10.1101/2021.02.10.430614

**Authors:** Michalis Mastri, Swathi Ramakrishnan, Shruti Shah, Ellen Karasik, Bryan M Gillard, Michael M Moser, Bailey K Farmer, Gissou Azabdaftari, Gurkamal S Chatta, Anna Woloszynska, Kevin H Eng, Barbara A Foster, Wendy J Huss

## Abstract

Patient derived models (PDMs) are a powerful tool to study preclinical responses. However, the benefits of each model have not been compared head-to-head when models are derived from the same surgical specimen. PDMs derived from surgical specimens were established as xenografts (PDX), organoids (PDO), and spheroids (PDS). PDMs were molecularly characterized by RNA sequencing. Differential gene expression was determined between the PDMs and surgical specimens. Surgical specimens had the most differentially expressed genes reflecting loss of immune and stromal compartments in PDMs. PDMs and surgical specimens were clustered using the Euclidian distance analysis to test model fidelity. PDMs upregulated a clear, patient-specific bladder cancer signal. Overall, the molecular profiles of PDXs were the most similar to the matching patient surgical specimen than the PDO and PDS from that patient. The epithelial mesenchymal transition (EMT) gene expression profile is maintained in the PDMs showing the persistence of EMT in both *in vivo* and *in vitro* model setting. The consensus molecular subtype was determined in order to compare PDMs to each other and their matching surgical specimen, and only surgical specimens with Basal/Squamous or Luminal Papillary molecular subtype established PDMs. Patient derived models reduce tumor heterogeneity and allow analysis of specific tumor compartments while maintaining the gene expression profile representative of the original tumor.

## 1. Introduction

### Bladder cancer is a significant public health problem

In the United States, approximately 80,470 new patients were diagnosed with bladder cancer with an estimated 17,670 deaths in 2019 (seer.cancer.gov). The majority of cases is non-muscle invasive and is treated by cystoscopic resection, with or without intravesical medical therapy instilled directly into the bladder. However, approximately a third of newly diagnosed bladder cancers have muscle-invasive bladder cancer (MIBC). Therapy for these patients consists of either a radical cystectomy or definitive chemoradiation therapy. Even with definitive treatment, the mortality from MIBC remains high.

### Consensus Molecular Classification of MIBC

Platinum-based chemotherapy continues to be the mainstay of systemic therapy for MIBC. With the exponential increase in knowledge about the molecular taxonomy of bladder cancer, additional therapeutic modalities are being rapidly developed. The recent development of a “consensus” classification system allows application of the molecular subtyping in the clinic to predict more effective treatment options.^1^ The six molecular subtypes in the consensus molecular classification based on 1750 MIBC from 6 datasets, arranged from most to least differentiated, are: Luminal Papillary (LumP) 24%, Luminal Non-Specified (LumNS) 8%, Luminal Unstable (LumU) 15%, Stroma-rich 15%, Basal/Squamous (Ba/Sq) 35%, and Neuroendocrine-like (NE-like) 3%.^1^ Although the overall survival outcome is improved with gemcitabine/cisplatin (G/C) neoadjuvant therapy the overall survival still remains poor.^2^ The LumP and Ba/Sq represent the 2 most common subtypes and account for 59% of MIBC cases. LumP and Ba/Sq subtypes are at opposite ends of the spectrum of differentiation. The median overall survival is highest for the more differentiated LumP at 4 years whereas the median overall survival for patients with the less differentiated Ba/Sq tumors is only 1.2 years.

Several studies, including the consensus molecular signatures, show that MIBC with a basal phenotype have the best outcomes with cisplatin-based neoadjuvant (NAC).^3, 4^ There are several weaknesses of molecular analysis in bulk tissue. With bulk RNA sequencing analysis it is unclear which cellular components are contributing to the signal since the sample tissue is comprised of multiple components including the tumor, stroma, and immune compartments. Thus, the selection process of PDM establishment may select for certain cell types and allow analysis of a more homogenous cell population. Integrating multi-omic analysis further predicted treatment response in bladder cancer^5^ and the use of PDMs can facilitate this analysis when patients’ specimens are limited or need to be reanalyzed with new technology.

## 2. Results

### 2.1. PDMs established from surgical specimens of bladder cancer

The ability of each tumor specimen to grow and establish a PDX model was determined by implanting 55 freshly procured bladder tumor specimens from TURBT or cystectomy procedures at Roswell Park. Nineteen (19) of 55 specimens grafted into host animals demonstrated tumor growth of at least 1 cm^3^ in the initial passage (34.5%; 95%CI: 21.6-47.5%). Initial growth of the grafted surgical specimen is considered p0. Nine (9) of the 19 tumors with initial growth (47.4%, 22.6-72.1%) established PDX tumor lines defined as established growth beyond p2 and demonstrated >80% tumor take rate after a viable freeze. Thus, 9 PDX tumor lines have been established from 55 patient specimens for a 16.4% (6.3-26.5%) take-rate, similar to previous studies.^6-9^ The histopathology, and deidentified demographic information of patient tumor specimens that resulted in established PDX models are listed in Table 1, and information for all patient tumor specimens tested for PDX establishment are listed in the Supplemental Table 1. There was no correlation between the ability to establish PDX models and tumor stage or treatment. Specimens were grafted and maintained in sex matched hosts. Interestingly, specimens from female patients were more likely to establish models at p0 compared to specimens from male patients (69.2% versus 23.8%, prop test p=0.0075), independent of tumor stage. PDOs and PDSs were established from either surgical specimens or xenografts to provide in vitro tools for future analysis; only RP-BL019 and RP-BL022 have a PDO and only RP-BL022 has a PDS derived from the surgical specimen. To determine whether gene expression changes are associated with specific PDMs, the RNA sequencing from each PDM was analyzed individually to identify differences within each patient surgical specimen and corresponding model.

**Table 1:** Surgical specimen demographics with established >p3 PDX models.

| RP-BL | Stage | Treatment | Response | Pathology | Sex | Age | Ethnicity | Smoking Status |
| --- | --- | --- | --- | --- | --- | --- | --- | --- |
| 003 | T3 | G/C | Non-Responder, Deceased | HG urothelial carcinoma | F | 57 | AA | Current |
| 005 | T2 | None | No Follow up | HG urothelial with sarcomatoid features | M | 83 | C | Former |
| 019 | T3 | G/C | Non-Responder, Deceased | HG urothelial carcinoma | F | 66 | C | Former |
| 022 | T2 | G/C | Non-Responder | HG papillary urothelial | M | 76 | C | Former |
| 040 | T1 | None | Deceased | HG with prominent squamous differentiation | F | 72 | C | Never |
| 050 | T3 | None | No Treatment | HG urothelial carcinoma with squamous differentiation | F | 76 | C | Never |
| 051 | T2 | None | Non-Responder, currently Pembrolizumab Treatment | HG papillary urothelial carcinoma | M | 60 | C | Never |
| 052* | T3 | BCG, G/C, pembro | Non-Responder, Deceased | Recurrence, HG papillary urothelial carcinoma | F | 64 | C | Former |
| 054 | T2 | None | No Treatment, Deceased | HG papillary urothelial carcinoma | F | 85 | C | Never |

\*Not sequenced due to multiple treatments of patient prior to specimen collection. BCG = Bacillus Calmette-Guerin; G/C = Gemcitabine/Cisplatin; Pembro=pembrolizumab; HG=high grade; F=Female; M=Male; C=Caucasian; AA=African American;

### 2.2. Molecular comparison of PDMs with original surgical specimen

#### 2.2.1. PDM transcriptomes faithfully represent their corresponding surgical specimen

The Euclidian distance between all the samples was calculated using the expression of highly expressed genes and specifically genes with higher expression in PDM compared to surgical specimens. These selection criteria filtered out genes that are underrepresented or not expressed in the tumor cells, like immune and stroma related genes. Using the Euclidian distance between the samples and the t-distributed stochastic neighboring embedding algorithm to visualize how similar the samples are, we found that samples cluster with their corresponding surgical specimen (Figure 1A). This clustering was persistent even when 3D cultures (defined as PDO and PDS) (Figure 1B), surgical specimens (Figure 1C), or surgical specimen and PDX (Figure 1D) were removed from the analysis reinforcing the finding that each PDM is more similar to its corresponding surgical specimen than its model type. Euclidean distance based on gene expression showed that PDX models are more similar to the surgical specimens followed by PDO and finally PDS (Figure 1E). For visualization, we identified differentially expressed genes by paired comparison of each PDM to its corresponding surgical specimen (Figure 1F) confirming that PDX are the most similar to surgical samples, whereas PDO and PDS are more similar to each other than surgical specimens or PDX. Of particular note, samples clustered based on patient or origin and not based on the model type, thus indicating that the PDMs are representative of its corresponding surgical specimen.

**Figure 1.**
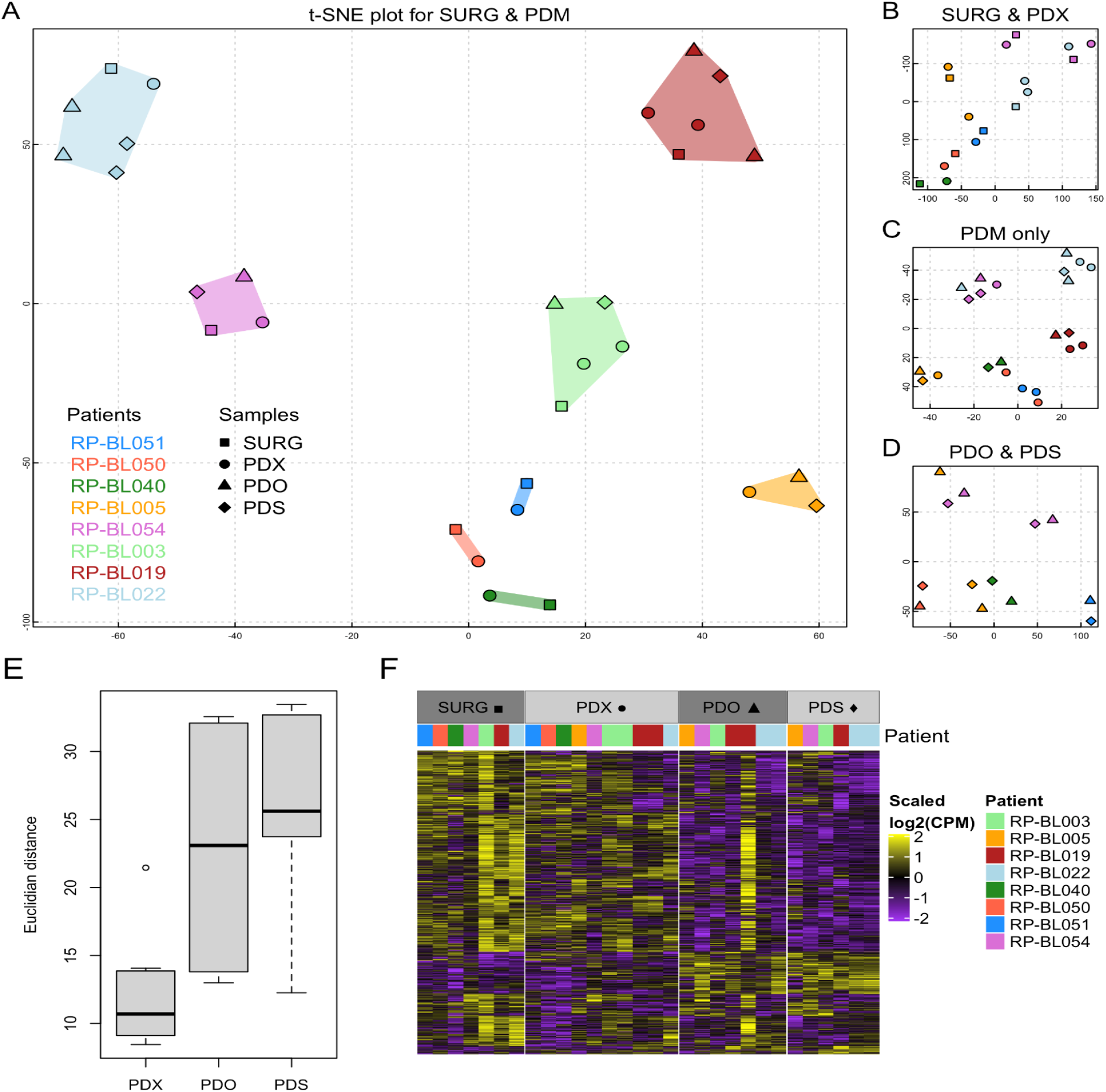
PDMs are a faithful representation of their original tumor. (A-D) Euclidean distance was visualized with t-SNE for (A) all samples, (B) surgical and PDX samples, (C) only PDM samples, and (D) PDO and PDS samples. (E) Euclidean distance of each PDM to their corresponding surgical specimen. (F) Heatmap of differentially expressed genes identified by paired comparison of PDM compared to their corresponding surgical specimen.

#### 2.2.2. PDMs amplify the tumor compartment transcriptome

To further examine the relationship between PDM and surgical specimen, differential gene expression was performed. Analysis between all PDMs and the surgical specimens showed that a plethora of genes are significantly downregulated in PDM (Figure 2A). We focused on 5 groups of genes related to tumor, bladder stem cells (BSD), general stem cells (SC), immune system, and the stroma. Only genes related to the immune system and the stroma were significantly downregulated in PDM, indicating that PDMs are representative of the tumor compartment but not the tumor microenvironment. Specific gene expression was visualized for each surgical specimen and PDM for the 5 tumor related groups of genes (Figure 2B). The gene expression pattern shows the expression of the individual genes in each of the samples emphasizing that the majority of the stroma and immune related genes are under-represented in PDM.

**Figure 2.**
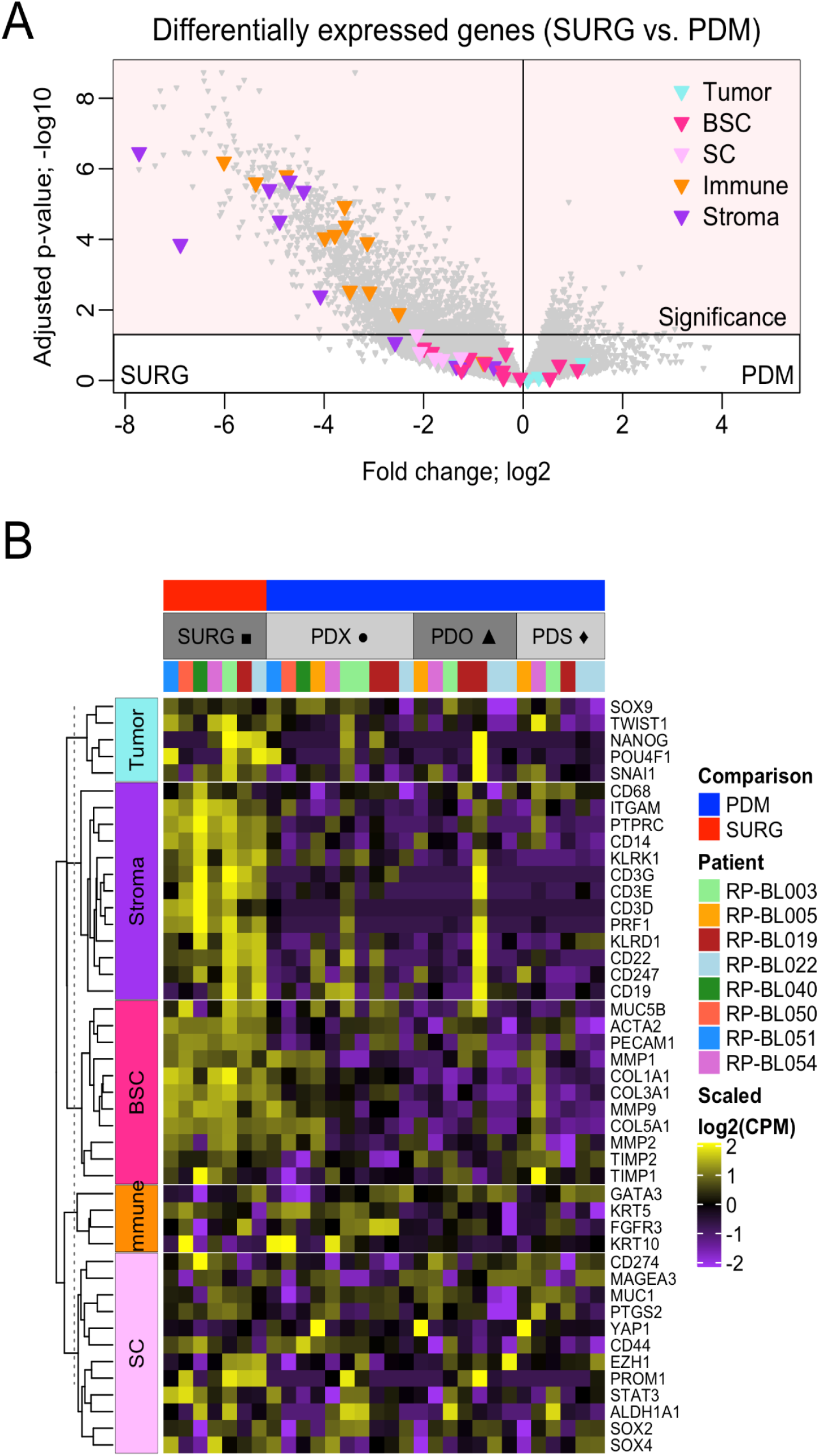
PDMs represent the tumor but not the TME. (A) Heatmap of gene expression corresponding to tumor, bladder cancer stem cell (BSC), stem cell (SC), immune, and stroma genes. (B) Volcano plots of differentially expressed genes comparing surgical specimens to PDM emphasizing the selected genes for tumor and tumor microenvironment.

#### 2.2.3. Hallmark gene set enrichment analysis in surgical specimens and PDMs

To further characterize the PDMs, gene set enrichment analysis for the hallmark gene sets was performed comparing surgical specimen vs. PDX, PDX vs. 3D models, and PDS vs. PDO (Figure 3A). The gene sets were grouped into 8 processes: development; DNA damage; immune; cellular component; metabolic; stress pathway; proliferation and signaling. Multiple gene sets were enriched in the surgical samples compared to the PDMs, but only the immune related gene sets were over-represented in the surgical specimens (Figure 3B). Interestingly, gene sets in the stress pathway group (apoptosis, protein secretion, reactive oxygen species, and unfolded protein) were enriched in the 3D culture PDM compared to the PDX (Figure 3C). Surprisingly, immune related gene sets were over-represented in PDS compared to PDO (Figure 3D) suggesting that immune related genes may be reactivated in spheroids due to the more stem-like phenotype of PDS model.

**Figure 3.**
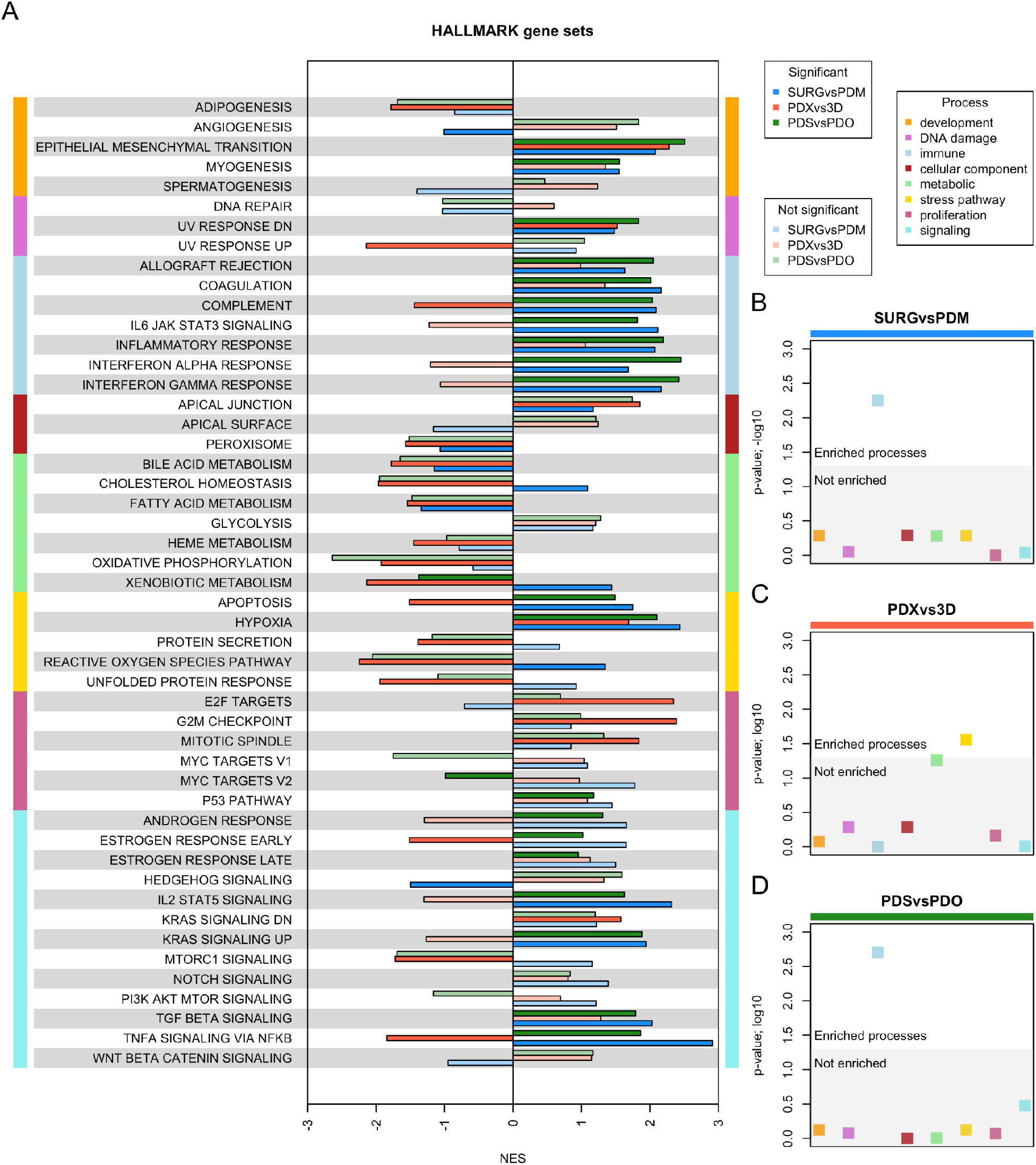
Hallmark pathway analysis RNAseq. (A) Gene set enrichment analysis for all the hallmark gene sets for 3 different comparison (SURG vs. PDM, PDX vs. 3D, PDS vs. PDO). Gene sets were categorized based on processes. (B-D) Enrichment of processes based on fisher exact test for (B) SURG vs PDM, (C) PDX vs 3D, and (D) PDS vs PDO.

#### 2.2.4. EMT and Molecular Subtypes of MIBC Analysis of PDMs and Surgical Specimens

The EMT hallmark gene set was enriched in all three comparisons (Figure 3A). The enrichment score for the EMT gene set showed that SURG is enriched vs. PDMs, PDX is enriched vs. 3D cultures and PDS is enriched vs. PDO models (Figure 4A). The EMT enrichment in PDS confirms the ability of cells with an EMT phenotype to establish as spheroids, compared to PDOs grown in media with differentiation factors. To determine how the EMT gene set was specific to the original surgical specimen derived models, Euclidean distance analysis was performed. The t-SNE representation of the Euclidean distance showed that expression of EMT genes was not enough to cluster the samples per patient, indicating that other biological processes define the surgical specimen (Figure 4B). The expression level of the EMT gene set for each surgical specimen and PDM shows that surgical specimens have increased levels of the EMT genes compared to the PDMs (Figure 4C). In addition, PDX models derived from patients that received treatment (RP-BL003, RP-BL019, and RP-BL022) had lower expression of EMT genes compared to the PDX derived from patients that were treatment naive. No pattern was obvious between PDO or PDS derived from patients that received treatment to those that were derived from patients that were treatment naive, indicating that the culture conditions used to derive the PDO and PDS alter the expression of the EMT genes.

**Figure 4.**
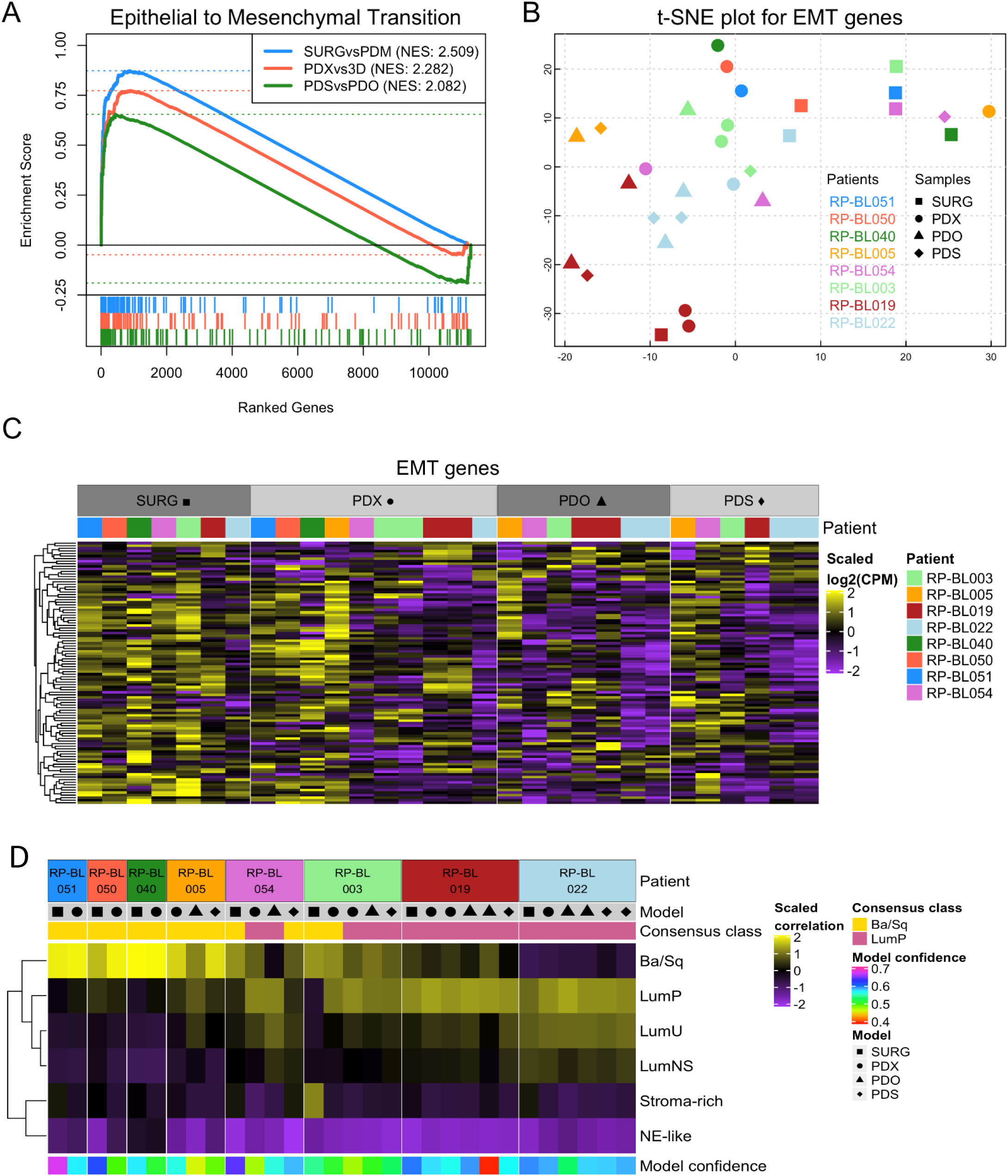
EMT and Ba/Sq molecular subtypes are enriched in PDMs. (A) Gene set enrichment analysis for the hallmark epithelial to mesenchymal transition gene set for 3 different comparisons (PDM vs. SURG, PDX vs. 3D, PDS vs. PDO). (B-C) Expression of the hallmark epithelial to mesenchymal transition genes were used for (B) t-SNE plot and (C) heatmap. (D) Correlation heatmap based on the consensus molecular subtype scores for each sample derived from RNA sequencing data.

The consensus molecular subtyping developed to classify MIBC^1^ was analyzed in order to determine if ETM maintenance in the PDMs was associated with any specific molecular subtype in the surgical specimens. The consensus molecular subtype classification of MIBC was analyzed for each surgical specimen and PDM to determine if the molecular subtype can be used to classify PDMs in order to provide pre-clinical models for each molecular subtype to facilitate treatment options. The molecular subtypes were determined for seven surgical specimens and their matched PDX models, as well as, six PDO and PDS models derived from the PDX models. Although the molecular subtype classification was designed for MIBC surgical specimens, the majority of the corresponding surgical specimens and their derived PDXs were classified as the same molecular subtype (Figure 4D). A high percent (5/7) of the surgical specimens that established PDMs were classified as the Basal/Squamous (Ba/Sq) molecular subtype, and most molecular classification of the PDMs corresponded to the surgical specimen molecular subtype. Interestingly, the Ba/Sq specimens were able to give rise to both Ba/Sq and Luminal Papillary (LumP) models (RP-BL054 and RP-BL003), whereas LumP specimens only gave rise to LumP models (RP-BL019 and RP-BL022).

### 2.3. Histopathology and differentiation marker analysis

Historically, the tumor stage and histopathology has provided key information to determine the course of treatment, however there is little association of histopathology, expression of differentiation markers and molecular subtypes in PDMs. Characterizing each model for histopathology and molecular subtype will allow better model selection for pre-clinical trials. To determine how well the consensus molecular phenotype associates with tumor histopathology, the available surgical specimens and PDXs were evaluated by a GU pathologist. The molecular subtype and histopathology were consistent between 5/8 surgical specimens and their derived PDX models. There was a discrepancy between the pathology and molecular subtypes in RP-BL005, 003, 022 PDXs (Table 2).

**Table 2.** Histopathology and molecular subtypes of surgical specimens and PDMs.

| Histopathology |  |  | Molecular Subtype |  |  |  |
| --- | --- | --- | --- | --- | --- | --- |
| RP-BL Line | Surgical Specimen | PDX | Surgical Specimen | PDX | PDO | PDS |
| 051 | HG papillary urothelial carcinoma | HG papillary urothelial carcinoma with squamous diff | Ba/Sq | Ba/Sq | ND | ND |
| 050 | HG urothelial carcinoma with squamous differentiation | HG papillary urothelial carcinoma with squamous diff | Ba/Sq | Ba/Sq | ND | ND |
| 040 | HG with prominent squamous differentiation | HG papillary urothelial carcinoma with squamous diff | Ba/Sq | Ba/Sq | ND | ND |
| 005 | HG urothelial with sarcomatoid features | HG papillary urothelial carcinoma with sarcomatoid | Ba/Sq | Ba/Sq | Ba/Sq | Ba/Sq |
| 054 | HG papillary urothelial carcinoma | HG papillary urothelial carcinoma with squamous diff and sarcomatoid | Ba/Sq | Ba/Sq | LumP | Ba/Sq |
| 003 | HG urothelial carcinoma | HG papillary urothelial carcinoma with squamous diff | Ba/Sq | LumP | LumP* | LumP* |
| 019 | HG urothelial carcinoma | HG papillary urothelial carcinoma with predominately sarcomatoid | LumP | LumP | LumP | LumP |
| 022 | HG papillary urothelial | NE differentiation | LumP | LumP | LumP | LumP |

Generally, the PDX histopathology was in agreement with that of the surgical specimen, although there was a trend towards the PDX being more undifferentiated with a gain of the squamous phenotype. In order to further elucidate the differences in the histopathology and the molecular subtype in RP-BL005, 003, and 022, IHC for markers of differentiation was performed in PDX tumors. PDX sections were analyzed by IHC to identify E-cadherin (epithelial cells), CK5 (basal marker) and CK 20 (superficial/intermediate urothelium marker), vimentin (EMT) and synaptophysin (NE marker) expressing cells (Figure 5). The IHC analysis for BL-051, 050, 040, 054, 019 were as expected based on the hisotopathology and molecular subtypes (Data Not Shown). PDX models were characterized based on pathology, molecular subtype, gene expression, and IHC. The RP-BL005 PDX has a sarcomatoid pathology, a Ba/Sq molecular subtype classification with high expression of E-cadherin and CK5 protein expression, but also expresses an EMT markers gene expression profile (Figure 4B) and vimentin protein expression. The RP-BL003 has a squamous differentiation pathology, LumP molecular subtype classification, while high E-cadherin and CK5 protein expression that confirms the squamous differentiation histopathology, but the low expression of CK20 does not correspond to the LumP molecular subtype; additionally, there is high synaptophysin protein expression that does not correspond to histopathology or molecular subtyping. The RP-BL022 PDX has NE differentiation histopathology, but a LumP molecular subtype. The expression of E-cadherin and CK20 protein expression supports the LumP molecular subtype, while synaptophysin protein expression supports the NE histopathology. The differentiation marker expression can help characterize the PDX models when there is discrepancy between the histopathology and the molecular subtype classification, but cells express multiple differentiation markers.

**Figure 5.**
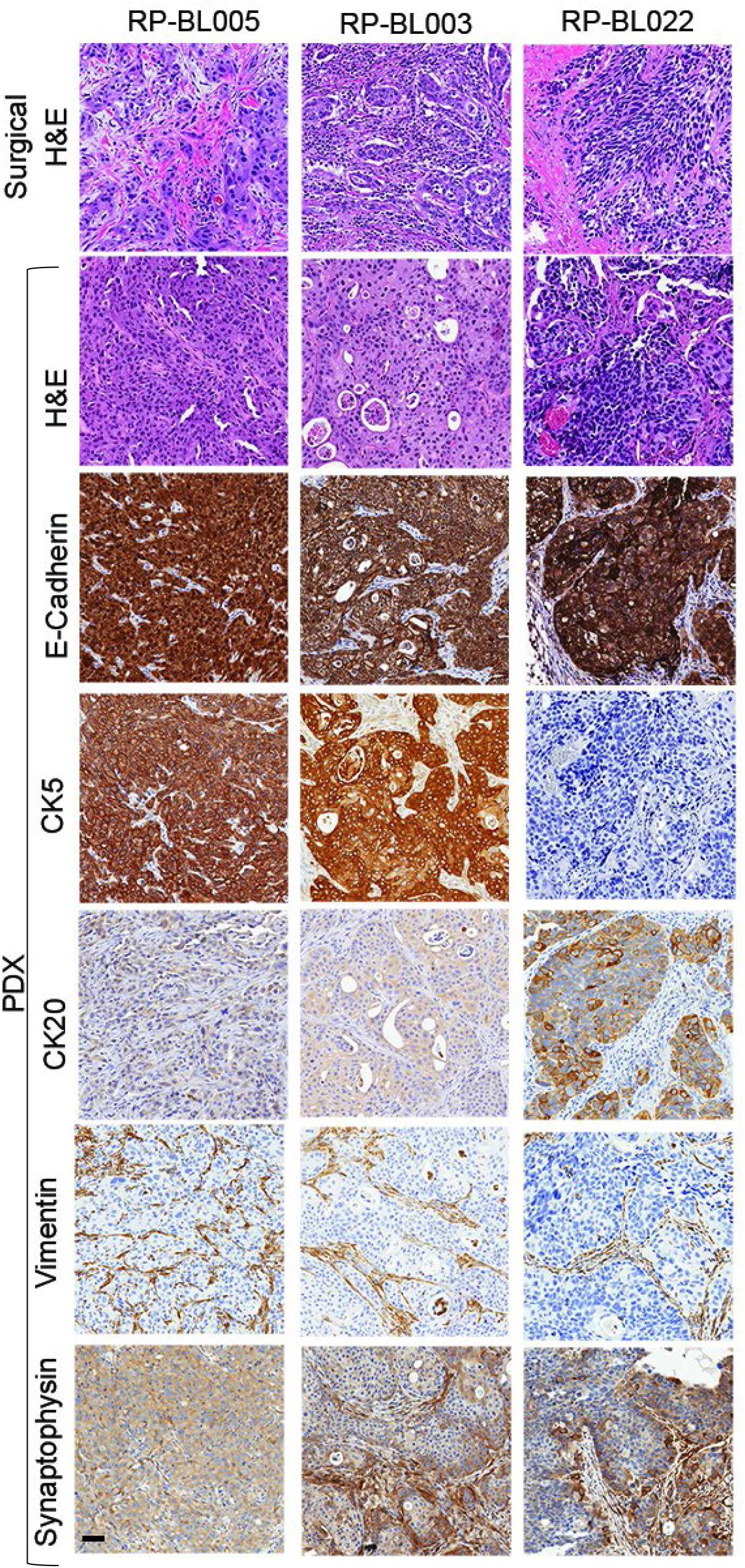
Pathology and differentiation marker analysis of PDMs with conflicting phenotypes. H&E staining of surgical specimens and PDX models derived from the surgical specimens PDX models RP-BL005, -003, and 022. IHC analysis of PDX models for E-Cadherin, CK5, CK20, Vimentin, and Synaptophysin. Bar = 50μm

## Discussion

The gene expression analysis comparing all three PDMs to the original surgical specimen allows the development of PDMs to test targeted and personalized therapy for bladder cancer. We classified the molecular subtype of 8 established bladder PDMs, and some PDMs may more faithfully reflect a particular molecular subtype, whereas others may better capture treatment response. In upper tract urothelial cancer PDX models, surgical specimens (growth and no growth n=70) were classified as primarily LumP (82.5%), LumU (8.75%), LumNS, Stroma-rich, and Ba/Sq (1.25%). There was a 16/17 histological concordance, and trend toward more invasive specimens more likely to establish PDX models.^10^ Our studies, did not include upper tract urothelial cancer and the enrichment of Ba/Sq (5/7) in our studies highlights the differences between these two types of cancers and possibly the variation in model development between the two groups. An understanding of how preclinical models with different molecular subtypes respond to different therapeutic approaches will allow for personalized medicine and improve therapeutic outcomes. Several studies, including the consensus molecular signatures, show that MIBC with a basal phenotype have the best outcomes with cisplatin-based neoadjuvant (NAC) therapy.^3, 4^ Thus determining the treatment response in these PDMs is an important future study.

Each PDM has selective pressures that are likely to affect the population of cells within the model. PDX models largely retain the same mutations and biological responses to therapies as observed in the surgical specimen from which they were derived.^8, 11, 12^ PDXs can be used to evaluate the biological response to therapeutic agents and molecular manipulations.^13-15^ PDX models, in which a patient’s tumor is grafted into an immunocompromised mouse, serve as a tool for preclinical investigation of tailored therapy and have the potential to meet the challenges described above. Advantages of PDXs include their ability to maintain original tumor heterogeneity and to circumvent confounding issues such as altered gene expression that result from serial passage of established cell lines grown on plastic. PDXs retain the variety of cell types found in the original tumor, such as vasculature, lymphatics, fibroblasts, smooth muscle, and, depending on the host, limited immune cells.^16^ Drawbacks of PDX models are contaminating mouse cells from the host, higher costs and longer time frames associated with working with mice compared to cells growing in culture, as well as the lack of an intact immune system in the host animal. Work by various investigators highlights the promise of personalized PDXs to predict drug response in the clinical setting, development of predictive biomarkers, and understanding mechanisms of treatment response or resistance.^8, 12, 17, 18^ PDXs offer a unique opportunity to evaluate the therapeutic response of a single tumor to multiple agents including growth of the tumor with no treatment.

The growth conditions for PDOs establishment promotes differentiation and give rise to the various different cell types within a tumor; thus, PDOs may better reflect the bulk tumor phenotype and therapeutic initial response of the cancer tissue from which they are derived, relative to other experimental models like established 2D cell lines. Conversely, the PDS assay is often used to enrich for the cancer stem cell population and test cancer stem cell properties *in vitro*. ^19-22^ Quantitation of cell viability, sphere number and size following treatment can provide a straightforward readout of therapy effectiveness on the cancer stem cell population. Cancer stem cells are speculated to be a potential source of therapy-resistant cells leading to recurrent disease.^23^ The PDS model is the most likely model to contain a high number of cancer stem cells and may have the least selective pressure to adapt to culture and the least heterogeneity because each sphere is clonally derived. Our data showed that PDS have a strong ETM gene expression profile compared to PDO (Figure 4A), confirming that PDS are more mesenchymal whereas PDO are more epithelial.

The PDMs were analyzed to determine if there was a common selection type for a particular PDM, were PDM more similar to each other or is the gene expression more like the original surgical specimen. Interestingly, all PDMs were more similar to the surgical specimen they were derived from (Figure 1). Transcriptomic data supports that variability is derived from interpatient differences and not the models. In addition, all PDMs amplify tumor signal, thus, we have a way to “purify” messy tumors before evaluating their transcriptome. Finally, PDMs reduce tumor heterogeneity and allow analysis of specific tumor compartments while maintaining the gene expression profile representative of the original tumor.

## 4. Materials and Methods

All materials used are provided in Supplemental Tables 2-5 and were sourced from companies in the USA unless indicated.

### 4.1 Ethics Statement

All of the tissue samples were collected under an Institutional Review Board (IRB)-approved protocol at Roswell Park Comprehensive Cancer Center. Specimens were collected after IRB-approved written consent from the patient was obtained at Roswell Park. All experiments were conducted and approved under our Institutional Animal Care and Use Committee (IACUC) protocol at Roswell Park.

### 4.2 Human Specimen Procurement

Fresh human bladder tissue, procured from transurethral resection of bladder tumor (TURBT) surgical specimens or radical cystectomies, were stored in static preservation solution (SPS-1™) up to 16 hours at 4°C. Tumor specimens were obtained from the Pathology Network Shared Resource at Roswell Park. Deidentified demographics (clinical stage, procedure, gender, age, ethnicity, smoking status, and treatment prior to specimen collection) are listed in Supplemental Table 1.

### 4.3 Patient Derived Models

#### 4.3.1 Xenograft Generation

The Experimental Tumor Model shared resource received patient samples for grafting into sex matched NOD.Cg-Prkdc^*scid*^ Il2rg^*tm1Wjl*^/SzJ mice (JAX, Bar Harbor, ME) hosts, also known as NOD SCID gamma (NSG). Patient samples that grow in NSG are designated as p0, and the next passage in NSG is p1, and so on. For bladder tumor samples, 0.5 -1 mm^3^ tumor pieces are dipped in Matrigel® and grafted subcutaneously (subQ) into the flank of host NSG mice. From the patient sample, 5-8 gender matched NSG hosts are grafted. PDX-p0s that grow from the patient sample are expanded by two additional rounds of growth (p1, p2) in sex matched NSG hosts.

#### 4.3.2 Organoid and Spheroid generation and culture

Tumor tissue from surgical specimens and PDX models was minced and enzymatically digested in a modified protocol from preciously published protocols.^24, 25^ Briefly, samples are incubated in a collagenase, dispase and DNAse solution (Supplemental Table 3) in a 50 mL flask for 1-2 hours at 37°C with gentle stirring. If necessary, red blood cells are removed by lysis using RBC lysis buffer, epithelial cells can be isolated with a Histopaque gradient and plated for 3D culture as previously described.^25^ Single cells are passed through a 100 µm cell strainer and resuspended in media appropriate for either organoid^26^ or spheroid^27, 28^ generation. Media components are listed in Supplemental Table 4.

For organoid growth, viable suspended cells were cultured in 5% Matrigel in defined organoid media mixture using the base R-spondin organoid culture system^26^ with growth factor conditions adapted for use with urothelial TCC. PDOs were expanded at a low rate of 2-4 fold after a dispase digestion (2.4 Units/ml in DPBS) of Matrigel followed by PDO digestion by TrypLE Express. Spheroids were established by seeding 1×10^3^ viable single cells/well in 24-well, ultra-low adhesion plates in 5% Matrigel in defined spheroid media. PDSs were passaged by digestion of the Matrigel with dispase, followed by digestion of the spheroids with trypsin. Once PDOs and PDSs reach p2 growth in multiple wells, aliquots were cryopreserved for RNA sequencing and a viable freeze in 10% DMSO.

### 4.4 Authentication of models and patient surgical samples

Short tandem repeats (STR) profiles were performed by Roswell Park’s Genomic Shared Resource (GSR) to authenticate that PDXs are derived from the matching patient sample. Flash frozen patient tumor samples were collected at the time of procurement and sent to GSR for DNA isolation and STR profile analysis. AmpFLSTR® Identifiler® Plus PCR Amplification Kit for STR profiling which utilizes fifteen STR loci and a sex-determining marker Amelogenin. PDX STR profile was compared to the patient tumor STR profile. A match of patient and PDM was called if the STR profile has a ≥90% match.

### 4.5 Histology and Immunohistochemisty (IHC)

PDX tissues were embedded in paraffin. Serial sections (5 μm) were cut and mounted on glass slides. Slides were deparaffinized in xylene, rehydrated through a graded series of alcohol washes, and equilibrated in double distilled water. Slides were incubated in 1x pH6 citrate buffer using DAKO PT link for 20 minutes, moved to DAKO Autostainer Plus for incubation in 3% H2O2 for 15 min. To block non-specific binding, tissues were incubated with 10% normal goat serum for 10 min, followed by avidin/biotin block. Antibodies used are listed in Supplemental Table 5. Primary antibodies E-Cadherin, CK5, CK20, Synaptophysin, Vimentin were diluted in 1% BSA solution and incubated for 30 minutes at room temperature All of the slides were incubated with the Goat anti Rabbit biotinylated secondary antibody for 15 minutes at room temperature. For signal enhancement, ABC reagent was applied for 30 minutes. To reveal endogenous peroxidase activity, slides were incubated with DAB substrate for 5 minutes and counterstained with DAKO Hematoxylin for 20 seconds.

### 4.6 RNA Isolation and Sequencing

RNA/DNA extraction was performed in the Genomics Shared Resource (GSR) at Roswell Park. The purification of total RNA is prepared using the miRNeasy micro kit (Qiagen 217084). Frozen tissues, organoid suspensions, and spheroid suspension samples are first suspended in 700 μl of Qiazol reagent. The samples were homogenized using Navy Rhino tubes in a Bullet Blender Homogenizer (Next Advance) for 5 minutes. The homogenate was removed and incubated in a new tube at room temperature. After addition of chloroform, the homogenate was separated into aqueous and organic phases by centrifugation. RNA partitions to the upper, aqueous phase, while DNA partitions to the interphase and proteins to the lower, organic phase or the interphase. The upper, aqueous phase is extracted, and ethanol is added to provide appropriate binding conditions for all RNA molecules from 18 nucleotides upwards. The sample is applied to the miRNeasy micro spin column, where the total RNA binds to the membrane and phenol and other contaminants are efficiently washed away. On-column DNAse digestion was performed to remove any residual genomic DNA contamination followed by additional washes. High quality RNA was eluted in 25 μl of RNase-free water. Quantitative assessment of the purified total RNA is accomplished by using a Qubit High Sensitivity RNA kit (Thermofisher), and concentration is determined by Ribogreen fluorescent binding to isolated RNA. The RNA was further evaluated qualitatively using RNA High Sensitivity tape on the 4200 Tapestation (Agilent technologies), where sizing of the RNA was determined, and a qualitative numerical score (RINe) is assigned. Amplified cDNA was generated using the SMART-Seq v4 Ultra Low Input RNA kit (Clonetech). 10 ng of total RNA was fragmented based on %DV200 analysis and used to synthesize first-strand cDNA utilizing proprietary template switching oligos. Amplified double strand (ds) cDNA was created by LD PCR using blocked PCR primers and unique sample barcodes are incorporated. The resulting ds cDNA was purified using AmpureXP beads (Beckman Coulter). Abundant Ribosomal cDNA was depleted using R probes, and 13 cycles of PCR using universal PCR primers to complete the library. The final libraries were purified using AmpureXP beads and validated for appropriate size on a 4200 TapeStation D1000 Screentape (Agilent Technologies, Inc.). The libraries are quantitated using KAPA Biosystems qPCR kit, and were pooled together in an equimolar fashion, following experimental design criteria. Each pool was denatured and diluted to 400pM with 1% PhiX control library. The resulting pool was loaded into 200cycle NovaSeq Reagent cartridge for 2×100 sequencing and sequenced on a NovaSeq6000 following the manufacturer’s recommended protocol (Illumina Inc.).

### 4.7 Bioinformatics

#### 4.7.1 RNA data processing

Sequencing quality control was assessed using FASTQC v0.11.5 (available online at: http://www.bioinformatics.babraham.ac.uk/projects/fastqc/). Reads were aligned to the human genome GRCh38 release 27 (Gencode) using STAR v2.6.0a^29^ and post-alignment quality control was assessed using RSeQC v2.6.5.^30^ Aligned reads were quantified at the gene level using RSEM v1.3.1.^31^ RSEM estimated gene counts were filtered and upper quartile normalized using the R-based Bioconductor package edgeR.^32^ Differential gene expression was performed after voom transformation followed by linear regression using the R-based Bioconductor package Limma.^33^

#### 4.7.2 Euclidean distance calculations

For t-distributed stochastic neighbor embedding (t-SNE) we first defined Euclidean distance based on the expression of a subset of genes. Genes were selected based on the average expression in all PDM and surgical specimens. Genes highly expressed in PDMs compared to surgical specimens were chosen. t-SNE plots were made using the R-based package Rtsne (https://github.com/jkrijthe/Rtsne).^34, 35 36^

#### 4.7.3 Hallmark Gene Set, EMT, consensus molecular subtype analysis

Gene set enrichment analysis was performed using the R-based Bioconductor package fgsea ^37^ for the hallmark gene sets from the molecular signatures database (MSigDB).^38, 39^ Molecular subtype classifications were performed on all surgical and PDM specimens using the consensus MIBC R package.^4^ RNA sequencing data were used and molecular subtype was assigned to samples only if it had a correlation value greater than 0.3. This package can identify 6 molecular classes: Luminal Papillary (LumP), Luminal Non Specified (LumNS), Luminal Unstable (LumU), Stroma-rich, Basal/Squamous (Ba/Sq), Neuroendocrine-like (NE-like).

## Supplementary Materials

Supplemental Tables 1-5

