## Supplemental Tables for "Patient derived models of bladder cancer amplify tumor specific gene expression compared to surgical specimen while maintaining gene expression of molecular subtype and epithelial mesenchymal transition markers"

S. Table 1. Patient demographics for all specimens where PDMs were attempted.

| RP-BL | Growth | Tumor Stage | Treatment | Pathology | Procedure | Sex | Age | Ethnicity | Smoking Status |
|---|---|---|---|---|---|---|---|---|---|
| 002 | No | T3 | None | HG Urothelial carcinoma with nested and lymphoepithelial features | TURBT | M | 75 | C | Never |
| 003 | Est | T3 | G/C | HG urothelial carcinoma | TURBT | F | 57 | AA | Current |
| 004 | No | T1 | BCG | HG urothelial | TURBT | M | 79 | C | Former |
| 005 | Est | T2 | None | HG urothelial with sarcomatoid features | TURBT | M | 83 | C | Former |
| 006 | No | T2 | BCG | Recurrrence HG urothelial carcinoma | TURBT | M | 55 | C | Current |
| 007 | No | T2 | BCG | HG papillary urothelial | Cystoprost atectomy | M | 84 | AA | Former |
| 008 | No | T1 | BCG | HG papillary urothelial | TURBT | M | 68 | AA | Current |
| 009 | Yes | T1 | None | HG papillary urothelial | TURBT | M | 58 | C | Never |
| 010 | Yes | T2 | None | HG urothelial carcinoma | TURBT | F | 64 | C | Current |
| 012 | Yes | T2 | None | HG urothelial carcinoma | TURBT | F | 65 | C | Current |
| 013 | No | T1 | BCG | HG papillary urothelial | TURBT | M | 66 | C | Never |
| 014 | No | T2 | None | HG urothelial carcinoma | TURBT | M | 81 | C | Former |
| 015 | No | T3 | Atezolizumab | HG urothelial carcinoma, plasmacytoid variant | Cystoprost atectomy | M | 72 | C | Current |
| 016 | No | T1 | Atezolizumab | HG papillary urothelial | TURBT | F | 85 | C | Former |
| 017 | No | T2 | None | HG papillary urothelial | TURBT | M | 67 | C | Former |
| 018 | No | T2 | Pembro | HG papillary urothelial | TURBT | M | 63 | C | Current |
| 019 | Est | T3 | G/C | HG urothelial carcinoma | Cystectomy | F | 66 | C | Former |
| 020 | No | T1 | None | HG papillary urothelial | TURBT | M | 68 | C | Former |
| 022 | Est | T2 | G/C | HG papillary urothelial | TURBT | M | 76 | C | Former |
| 024 | No | T2 | None | HG urothelial carcinoma with predominantly sarcomatoid | Cystoprost atectomy | M | 75 | C | Former |
| 026 | Yes | T4 | None | HG urothelial carcinoma | Cystectomy | F | 69 | C | Current |
| 027 | No | T4 | None | HG urothelial carcinoma | Cystoprost atectomy | M | 72 | C | Former |
| 028 | No | Met | Multiple Chemo | Met-Poorly diff | TURBT | M | 70 | C | Never |
| 029 | Yes | T1 | BCG+INF | HG papillary urothelial | Cystoprost atectomy | M | 46 | C | Current |
| 030 | Yes | T1 | Treatment for Multiple Myeloma* | HG urothelial carcinoma | TURBT | M | 83 | C | Former |
| 031 | No | T1 | None | HG papillary urothelial | Cystoprost atectomy | M | 84 | C | Never |
| 032 | No | T1 | None | HG urothelial carcinoma with squamous differentiation | TURBT | M | 86 | C | Never |
| 033 | No | T1 | None | HG urothelial | TURBT | M | 52 | C | Former |

| | | | | | | | | | |
|---|---|---|---|---|---|---|---|---|---|
| 034 | No | T4 | None | HG urothelial carcinoma with focal glandular and squamous differentiation | TURBT | M | 64 | C | Never |
| 036 | Yes | T1 | None | HG papillary urothelial | TURBT | M | 76 | C | Former |
| 037 | Yes | T1 | None | HG papillary urothelial | TURBT | M | 52 | C | Former |
| 038 | No | T1 | None | HG papillary carcinoma | TURBT | M | 75 | C | Former |
| 039 | No | T1 | None | HG papillary urothelial | TURBT | M | 76 | C | Current |
| 040 | Est | T1 | None | HG with prominent squamous differentiation | TURBT | F | 72 | C | Never |
| 041 | No | T2 | None | HG papillary urothelial | TURBT | M | 90 | C | Former |
| 042 | No | T2 | G/C | HG papillary urothelial carcinoma with focal squamous differentiation | TURBT | M | 76 | C | Never |
| 043 | No | T2 | None | Poorly differentiated urothelial carcinoma w/ basal-squamous differentiation | TURBT | M | 47 | C | Former |
| 044 | No | Ta | None | HG papillary urothelial | TURBT | M | 68 | C | Current |
| 046 | No | T1 | None | HG urothelial carcinoma | TURBT | M | 54 | C | Current |
| 047 | No | T1 | None | HG papillary urothelial | TURBT | F | 82 | C | Former |
| 048 | No | T2 | None | HG papillary urothelial carcinoma | Cystoprostatectomy | M | 51 | C | Current |
| 049 | No | T4 | Pembro | HG papillary urothelial carcinoma | TURBT | M | 79 | C | Former |
| 050 | Est | T3 | None | HG urothelial carcinoma with squamous differentiation | TURBT | F | 76 | C | Never |
| 052 | Est | T3 | BCG, G/C, pembro | Recurrence, HG papillary urothelial carcinoma | TURBT | F | 64 | C | Former |
| 054 | Est | T2 | None | HG papillary urothelial carcinoma | TURBT | F | 85 | C | Never |
| 055 | NA | T2 | None | HG urothelial carcinoma w/ squamous diff | Cystoprostatectomy | F | 75 | C | Current |
| 056 | No | T3 | None | HG urothelial carcinoma w/ squamous diff | TURBT | F | 73 | C | Former |

BCG = Bacillus Calmette-Guerin; G/C = Gemcitabine/Cisplatin; INF = Interferon; Pembro=pembrolizumab; HG=high grade; TURBT=Transurethreal Resection Bladder tumor; F=Female; M=Male; C=Caucasian; AA=African American; NA=Not available. Growth column: Est=Established PDX model; Yes=Grew but did not establish passable PDX model; No=No Growth. Row shading indicates surgical specimen that established PDXs and are also in Table 1.

S. Table 2. Materials used

| Material | Source | Product Number |
|---|---|---|
| Histopaque | Sigma | 10771-500ml |
| Cell strainer | ThermoFisher Scientific | 352360 |
| Matrigel | Corning | 354234 |
| TrypLE Express | ThermoFisher Scientific | 12604013 |
| eBioscience™ 1x RBC Lysis Buffer | ThermoFisher Scientific | 00-4333-57 |
| 24-well, ultra-low adhesion plates | VWR | 734-2779 |
| DMSO | ThermoFisher Scientific | 50-980-367 |
| AmpFLSTR® Identifiler® Plus PCR Amplification Kit | ThermoFisher Scientific | A26364 |
| Fisherbrand probe on plus | Fisher Scientific | 22-230-900 |
| StarFrost Slides Adhesive | Mercedes Medical | MER7255/90/WH |
| Citrate Buffer | Invitrogen | 00-5000 |
| PBS | Invitrogen | 14190-144 |
| Normal goat serum | Invitrogen | 50062Z |
| Avidin/biotin block | Vector Labs | SP-2001 |
| BSA | ThermoFisher Scientific | BP1605-100 |
| ABC reagent | Vector Labs | PK 6100 |
| DAB substrate | DAKO | K3467 |
| Hematoxylin | DAKO | CS7000 |
| Belzer UW® Cold Storage Solution | Bridge to Life | NA |

S. Table 3. Dispase digestion solution materials

| Material | Source | Product number | Final Conc. |
|---|---|---|---|
| Deoxyribonuclease I | Sigma | DN25-1G | 0.01% |
| Dispase | ThermoFisher Scientific | 17105-04 | 2.4 Units/ml |
| Collagenase II | Sigma | C6885-1G | 0.28% |
| DPBS | Invitrogen | 14190-144 | |

S. Table 4. Spheroid and organoid media components

| Material | Source | Product number | PDS conc. | PDO conc. |
|---|---|---|---|---|
| Base Media | Invitrogen | 12637-010 | - | Advanced DMEM |
| | Invitrogen | 11320-033 | DMEM/F12 | - |
| B27 | Invitrogen | 17504-044 | 2× diluted | 50× diluted |
| *N*-acetylcysteine | Sigma | A9165 | - | 1.25 mM |
| EGF | Peprotech | AF-100-15 | 10 ng/ml | 5 ng/ml |
| Noggin | Peprotech | 120-10C | - | 100 ng/ml |
| R-spondin 1 | R&D Systems | 4645-RS-025 | - | 500 ng/ml or 10% conditioned medium |
| A83-01 | Tocris Bioscience | 2939 | - | 500 nM |
| FGF10 | Peprotech | 100-26 | - | 10 ng/ml |
| FGF2 | Peprotech | 100-18B | 5ng/ml | 5 ng/ml |
| Prostaglandin E2 | Tocris Bioscience | 2296 | - | 1 µM |
| Nicotinamide | Sigma | N0636 | - | 10 mM |
| SB202190 | Sigma | S7076 | - | 10 µM |
| Insulin | Invitrogen | 12585-014 | - | 4mg/ml |
| Y-27632 dihydrochloridea | Selleck Chemical | S1049 | - | 10 µM |

S. Table 5. Antibodies used for IHC analysis

| Antibody name | Source | Catalog Number | Concentration/Dilution |
|---|---|---|---|
| E-Cadherin | BD | 610181 | 1:500 |
| CK5 | Covance | PRB 160p-100 | 1:1000 |
| CK20 | Abcam | ab97511 | 1:500 |
| Synaptophysin | Abcam | ab52636 | 1:400 |
| Vimentin | Cell Signaling | 5741 | 1:400 |
| Goat anti Rabbit | Vector Biolabs | BA1000 | 1:600 |
